# TNFα hinders FGF4 efficacy to mitigate ALS astrocyte dysfunction and cGAS-STING pathway-induced innate immune reactivity

**DOI:** 10.1101/2023.11.08.566131

**Authors:** Erika Velasquez, Ekaterina Savchenko, Sara Marmolejo-Martínez-Artesero, Désiré Challuau, Aline Aebi, Yuriy Pomeshchik, Nuno Jorge Lamas, Mauno Vihinen, Melinda Rezeli, Bernard Schneider, Cedric Raoul, Laurent Roybon

## Abstract

Astrocytes play an important role in the onset and progression of amyotrophic lateral sclerosis (ALS), a fatal disorder characterized by the relentless degeneration of motor neurons (MNs) in the central nervous system. Despite evidence showing that ALS astrocytes are toxic to MNs, little is understood about the earliest pathological changes that lead to their neurotoxic phenotype. In this study, we generated human astrocytes from induced pluripotent stem cells (iPSCs) harboring the ALS-associated A4V mutation in superoxide dismutase 1 (SOD1), to examine cellular pathways and network changes similar to early stages of the disease. By using proteomics as a molecular indicator, we observed significant alterations in the levels of proteins linked to ALS pathology and the cGAS-STING pathway-induced innate immunity. Interestingly, we found that the protein profile of reactive ALS astrocytes differed from that of wildtype astrocytes treated with the pro-inflammatory cytokine TNFα. Notably, we showed that fibroblast growth factor 4 (FGF4) reversed ALS astrocyte dysfunction and reactivity, but failed to provide protection to MNs when expressed in the spinal cord of the SOD1^G93A^ mouse model of ALS. Further analysis showed that ALS astrocyte reactivity which was rescued by FGF4 was abrogated by TNFα. The latter is capable of exacerbating the dysfunction and reactivity of ALS astrocytes compared to control. Our data show that iPSC-derived ALS astrocytes are dysfunctional and spontaneously exhibit a reactive phenotype when generated from iPSCs. This suggests that this phenotype may resemble the early stages of the disease. Our data also demonstrate that reducing mutant astrocyte reactivity in vivo using FGF4 is not sufficient to prevent MN death in a mouse model of ALS. To mitigate ALS, future studies should investigate whether dual therapies that both lower astrocyte reactivity and reverse disease-associated cellular dysfunction could prevent MN death.

**Graphic abstract:** 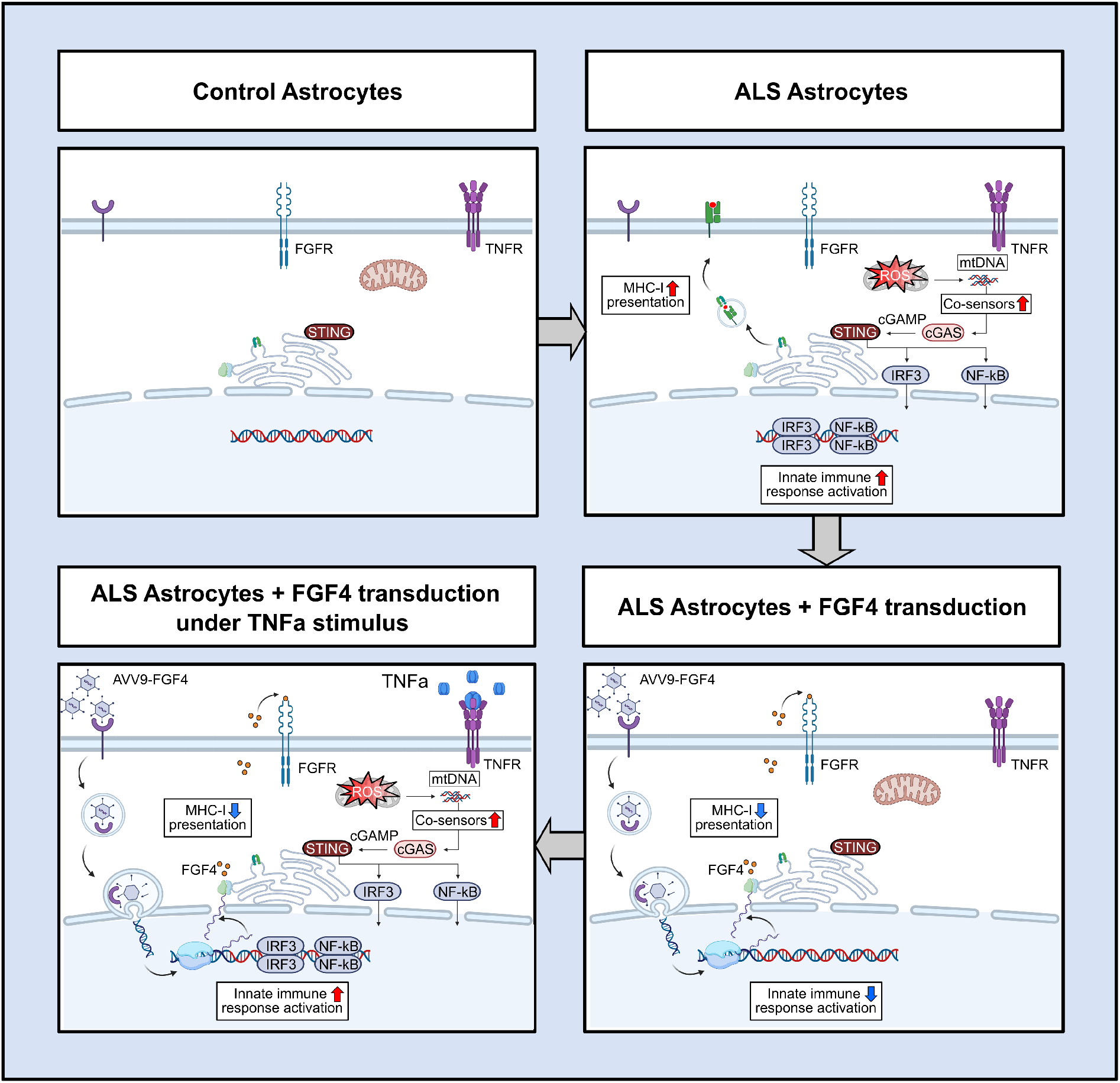

**Highlights:**

– ALS astrocytes are dysfunctional and reactive compared to wildtype astrocytes
– FGF4 reverses ALS astrocyte dysfunction and reactivity
– FGF4 lowers ALS astrocyte reactivity in vivo but fails to protect ALS motor neurons from death
– ALS astrocyte reactivity rescued by FGF4 is attenuated by TNFα

## Background

Amyotrophic lateral sclerosis (ALS) is a fatal disorder characterized by the progressive degeneration of motor neurons (MNs) in the motor cortex, brainstem and spinal cord (SC) [1, 2]. ALS is one of the most devastating adult-onset neurodegenerative diseases, with an estimated mortality of 25,000 patients yearly in the United States [3]. The disease leads to muscular weakness, atrophy and paralysis and ultimately causes death due to respiratory failure. Most ALS cases are sporadic (sALS), but approximately 5–8% are familial (fALS), with 15–20% harboring a mutation in the *superoxide dismutase 1* (*SOD1*) gene [4].

The etiology of ALS remains unknown. Accumulated data suggest that glial cells are critical for the onset and progression of the disease [5]. The selective expression of mutant SOD1 in MNs fails to induce neurodegeneration in mouse models, indicating an active involvement of neighboring cells in this process [6]. In ALS, astrocytes display a neurotoxic phenotype associated with the loss of their neuro-supportive functions [7–9]. Deficits in glutamate uptake by astrocytes have also been observed in ALS [10], with increased neurotransmitter accumulation in the synaptic cleft [11]. Mutant SOD1 astrocytes modulate the GluA2 subunit of AMPA receptors in MNs, which may confer vulnerability to glutamate excitotoxicity [12]. The presence of reactive astrocytes has been observed in the cortex and SC of ALS patients [13], releasing inflammatory and pro-apoptotic mediators that render MNs more susceptible to degeneration. The transplantation of mutant SOD1^G93A^ glial-restricted precursors and human sporadic ALS spinal neural progenitor cells in the cervical SC of wildtype (WT) mice induces MN degeneration *in vivo* [14], thus confirming the pivotal role of astrocytes in the pathogenic process underlying MN death. Despite all the evidence to date, the earliest pathological changes driving astrocyte dysfunction in ALS have not been fully elucidated.

One of the challenges in understanding the earliest molecular changes behind ALS progression is the limited access to brain samples from patients in the early stages of the disease, as well as the difficulty of determining the specific contribution of each cell type in complex tissues like the brain and SC. However, these limitations can be overcome by using patient-derived induced pluripotent stem cells (iPSCs). iPSC-derived astrocytes exhibit cellular pathogenic features that primarily reflect the activity of pathogenic genes or risk factors carried by the donor patient during prodromal disease stages. Moreover, this tool also represents a platform to genetically modulate astrocyte biology and modify astrocytes pathologic profile and toxicity to MNs [15–18].

Currently, there is no effective therapy for ALS. The available pharmacological options (such as riluzole [19] or edaravone [20]), together with palliative measures, only modestly increase the life expectancy of the patients. In this context, gene therapy targeting ALS astrocytes is a potential avenue to modulate their inflammatory profile and restore their neurotrophic support. Previous studies showed that healthy astrocytes can positively impact the disease course. Focal transplantation of WT glial-restricted precursors in the cervical SC of SOD1^G93A^ mice decreases MN loss and significantly extends animal survival [21].

Previously, we demonstrated the impact of fibroblast growth factor 2 (FGF2) treatment in WT astrocytes. FGF2 strongly increased astrocytic expression of the glutamate transporter- 1 (GLT1) [22, 23]. It has also been reported that FGF signaling in the neocortex is necessary to maintain the non-reactive state of astrocytes and accelerate their deactivation after injury [24]. All of this data suggests that the FGF pathway in astrocytes is a promising therapeutic candidate to be explored in ALS. Here, using astrocytes derived from iPSCs generated from patients harboring mutant SOD1^A4V^, we investigated the earliest protein changes related to ALS progression and the effect of adeno-associated virus (AAV)-mediated expression of several FGFs family members on the astrocyte proteome. We found significant alterations in the levels of proteins associated with the innate immune system which serve as early molecular signatures of ALS astrocytes. Furthermore, we discovered that FGF4 had the most substantial effect in counteracting the perturbation in the innate immune system *in vitro*. However, when, we delivered FGF4 using AAV serotype 9 in SOD1^G93A^ mice, it did not confer MN protection. Nonetheless, FGF4 significantly reduced the reactivity of spinal astrocytes in ALS mice.

## Results

### ALS astrocytes are dysfunctional and reactive compared to wildtype astrocytes

We first examined whether ALS astrocytes exhibit cell-autonomous dysfunction. We generated astrocytes from iPSCs obtained from ALS patients and healthy individuals. The ALS astrocytes carried the A4V mutation in the SOD1 protein, which is one of the most aggressive variations reducing the mean survival time to under 2 years after diagnosis [25]. After 100 days of differentiation, the astrocytes expressed canonical markers glial fibrillary acidic protein (GFAP), S100 calcium binding protein beta (S100b), Ezrin, glutamine synthetase (GS) and inhibitor of DNA binding 3 (ID3). There were no significant changes observed in the percentage of astrocytic markers between control and ALS astrocytes (Figure 1a and b). However, ALS astrocytes displayed signs of reactivity featured by increased GFAP intensity staining and the presence of multiple short cellular processes (Figure 1c). For proteomic analysis, 100-days-old astrocytes were seeded and maintained in growth medium for 5 days (D5) and 14 days (D14).

**Figure 1.**
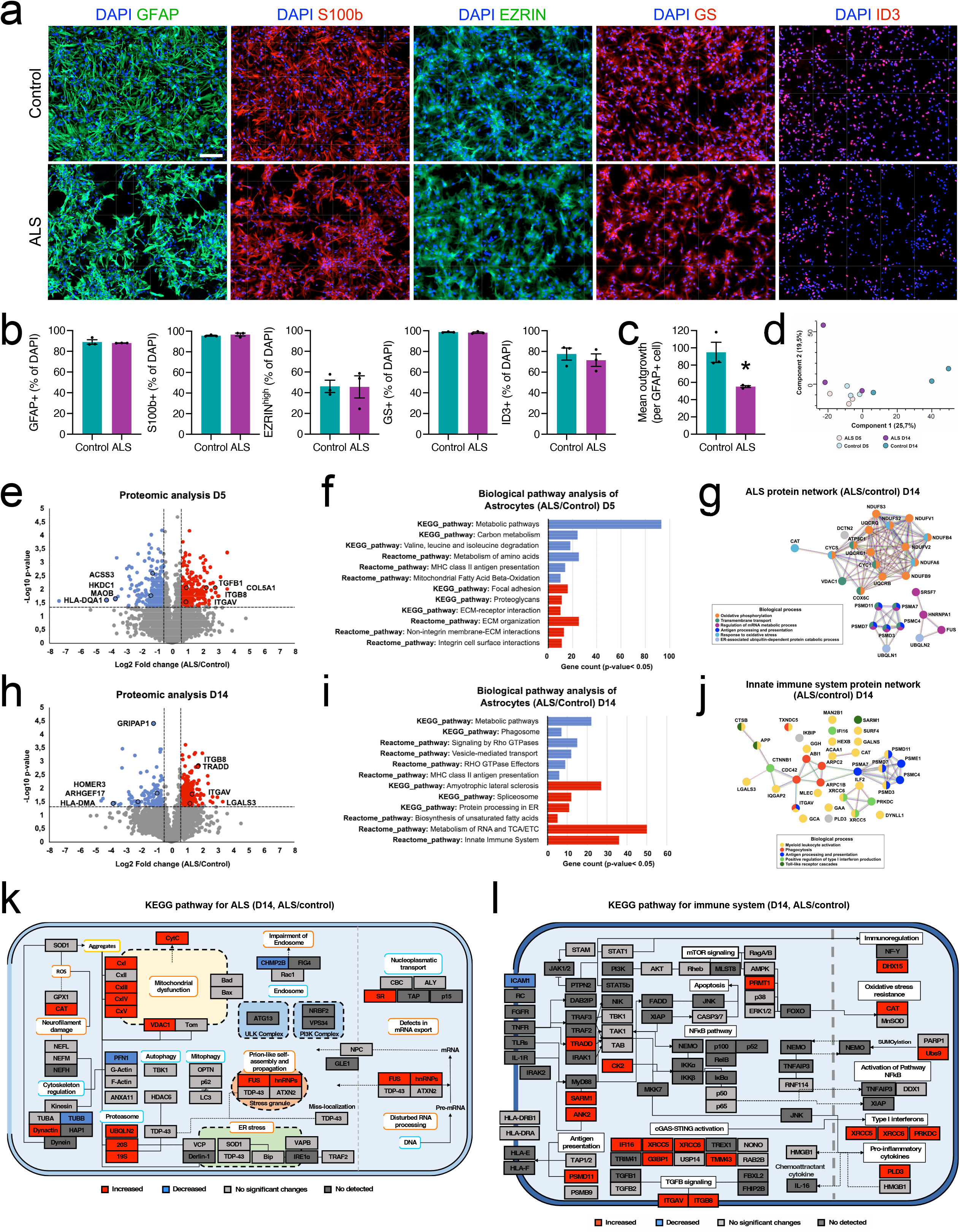
**a)** Representative immunostaining of cultures for canonical astrocytic markers glial fibrillary acidic protein (GFAP), S100 calcium-binding protein beta (S100b), membrane and actin cytoskeleton linker EZRIN, glutamine synthetase (GS) and inhibitor of DNA binding 3 (ID3) at day 100. Scale bar, 100 μm. **b)** Quantification of astrocytic markers in 100-day-old cultures generated from control and ALS iPSCs. Data are mean ± SEM; n = 3. **c)** Automated quantification of GFAP-positive cells with significant process outgrowth using the neurite outgrowth module of MetaMorph software. **d)** Principal component analysis (PCA) of astrocytes proteomes at 5 and 14 days. The PCA of the normalized protein intensities shows a separation of the protein profiles between controls and ALS cells from day 14. **e)** Volcano plot representing the proteins of ALS vs. control astrocytes at 5 days. Decreased and increased proteins are represented in blue and red, respectively (Two-tailed t-test p-value: 0.05; log2 fold-change cutoffs of + 0.5). **f)** Biological pathway analysis of dysregulated proteins between ALS and control astrocytes at 5 days (Enrichment p-value < 0.05). Pathways derived from down and up-regulated proteins are represented in blue and red, respectively. Detriment in the metabolic pathways, including amino acids and lipids, are the main alterations observed concomitantly with the increment of proteins related to the extracellular matrix. **g)** STRING interaction network analysis of the biological process of proteins associated with the ALS pathway altered in ALS astrocytes. **h)** Volcano plot representation of the proteome profile of ALS vs. control astrocytes at 14 days. Two-tailed t-test p-value: 0.05; log2 fold-change cutoffs of + 0.5. Decreased and increased proteins are represented in blue and red, respectively. **i)** Biological pathway analysis of dysregulated proteins in ALS vs. control astrocytes at 14 days (Enrichment p-value < 0.05). Pathways derived from down and up-regulated proteins are represented in blue and red, respectively. The decrease of proteins in metabolic and vesicle- related pathways and the increase of proteins linked to the immune system, RNA, and TCA/ETC functions, are the main findings on day 14. Additionally, from day 14, the enrichment of proteins previously associated with ALS disease starts to rise. **j)** STRING interaction network analysis of the biological process of proteins related to the innate immune system pathway altered in ALS astrocytes. **k)** ALS mapping based on KEGG mapper color analysis. Protein abundances of ALS vs. control astrocytes at 14 days were considered in this representation (Two-tailed t-test p-value: 0.05; log2 fold-change cutoffs of + 0.5). The ALS pathway map reveals profound alterations in the mitochondrial respiratory chain complexes and the proteasome system. **l)** Immune mapping based on KEGG mapper color and Reactome analysis. Protein abundances of ALS vs. control astrocytes at 14 days were considered in this representation (Two-tailed t-test p-value: 0.05; log2 fold-change cutoffs of + 0.5). The signaling map displays a basal activation of the innate immune system in the ALS astrocytes.

Mass spectrometry analysis of human astrocytes at 5 and 14 days of culture identified 7541 proteins in the dataset (Table S1). PCA analysis showed a clear separation in the proteome profiles of control and ALS astrocytes at D5 and D14 (Figure 1d). At D5, ALS astrocytes exhibited a decrease in the abundance of proteins related to metabolic pathways, such as amino acid and fatty acid metabolism, along with an increase in the expression of proteins associated with the extracellular matrix (ECM) (Figure 1e and f). The decreased protein expression in metabolic pathways persisted in ALS astrocytes at D14, with additional disruption in phagosome and vesicle transport proteins (Figure 1h and i). Furthermore, at D14, we observed an increase in the levels of proteins connected to ALS pathways, RNA and tricarboxylic acid cycle/electron transport chain (TCA/ETC) metabolism, ER protein processing, and the innate immune system (Figure 1h and i).

We the conducted a protein interaction network analysis to extract the main hubs and dissect the primary biological processes involved in the ALS and innate immune pathways. We found that ALS protein network encompassed oxidative phosphorylation, mRNA metabolism regulation and oxidative stress response (Figure 1g). In contrast, the innate immune system network predominated the myeloid leukocyte activation, including the positive regulation of type I interferon production, phagocytosis, and antigen presentation (Figure 1j).

Finally, we constructed a basic cell signalling map of ALS and immune system pathways based on the KEGG mapper to pinpoint the main proteins leading to these pathological changes. ALS mapping (Figure 1k) revealed perturbations in mitochondrial proteins, with increased levels of almost all components of the respiratory complex (CxI, CxII, CxIV, CxV), voltage-dependent anion-selective channel 1 (VDAC1), and cytochrome C. Proteins linked to ubiquitin-proteasome system (e.g., UBQLN2) were also altered in parallel with the increment of proteins pathologically related to ALS such as FUS [26, 27] and hnRNPs [28]. The immune system mapping (Figure 1l) showed the increase of several proteins related to innate immune response through the cyclic GMP-AMP synthase - Stimulator of interferon genes (cGAS-STING) pathway [29] (e.g., XRCC5, XRCC6, TMEM43) and transforming growth factor beta-1 activation (i.e., ITGAV, ITGB8).

Together, these data indicate that ALS astrocytes display an aberrant proteome profile with a higher basal protein expression. This higher expression then triggers mitochondrial and proteasomal dysfunction, along with the activation of innate immunity.

### ALS astrocyte reactivity differs from TNFα-induced astrocyte reactivity

Given that ALS-astrocytes have a more pronounced pro-inflammatory profile, we asked whether challenging WT astrocytes with tumor necrosis factor-alpha (TNFα) would result in the same proteome alterations observed in mutant SOD1 astrocytes. TNFα is a cytokine that is up-regulated from pre-symptomatic stage to the end stage in ALS patients and in animal models of the disease [30]. First, we separately evaluated the effect of TNFα treatment in control and SOD1^A4V^ astrocytes (Table S2). As expected, TNFα induced an increase in crucial inflammation-related proteins (e.g., TAP1/2, ICAM1) in both conditions (Figure 2a). However, in ALS astrocytes, we identified a 2- to 3-fold higher response to TNFα in signalling pathways related to antigen presentation and nuclear factor kappa beta (NF-κB) (Figure 2a). Pathway analysis in the control condition revealed a reduction in lysosome, glycan, and lipid metabolism and an increase in proteins involved in cytokine signalling, ER-phagosome pathway, and proteasome (Figure 2b). In ALS condition, membrane traffic, mTOR signalling, and lysosomal/peroxisomal metabolism were the main pathways downregulation, while the immune system, cytokine and focal adhesion pathways were upregulated (Figure 2b).

**Figure 2.**
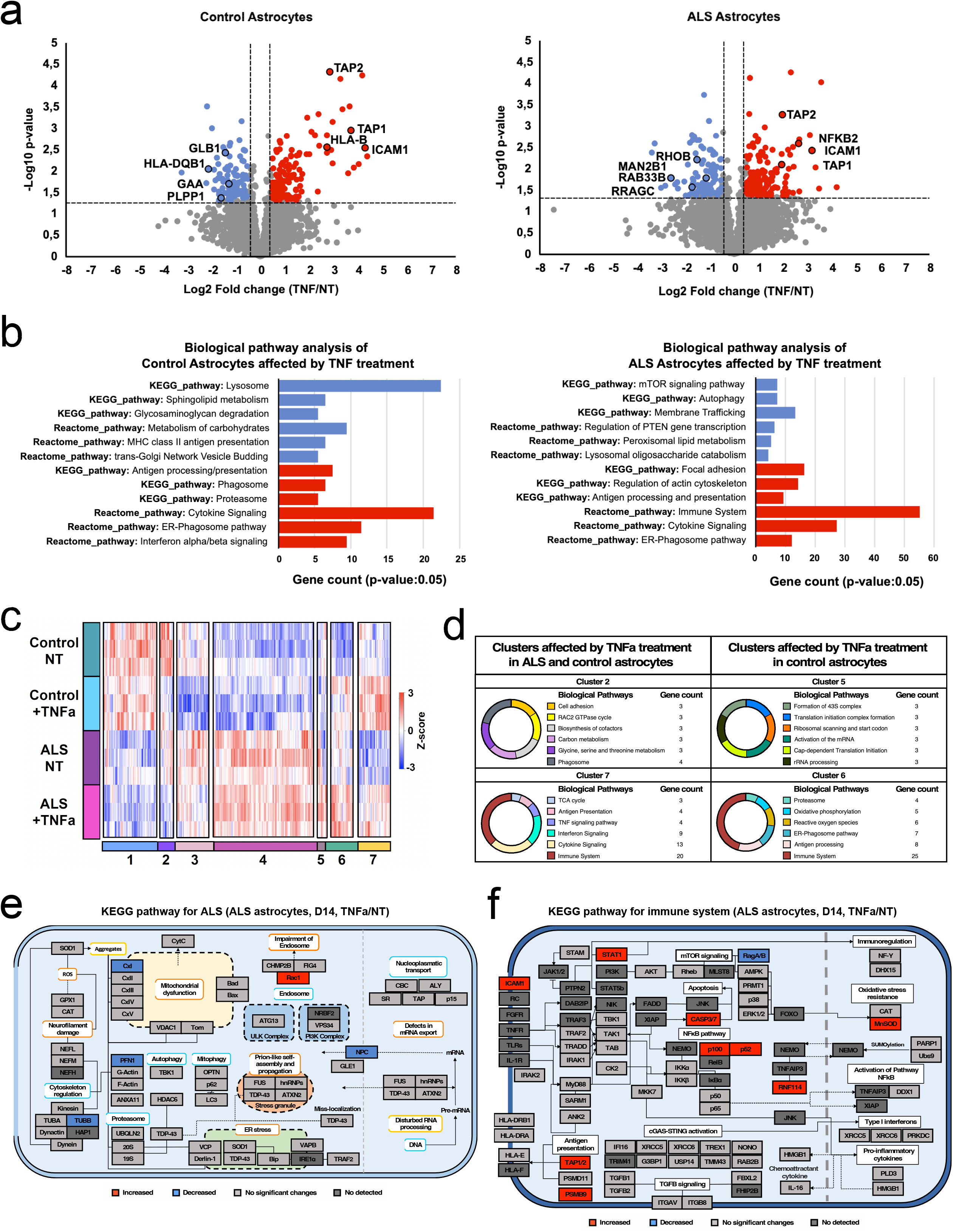
**a)** Volcano plot of the proteome profile of control and ALS astrocytes at 14 days treated with tumor necrosis factor-alpha (TNF) and not treated (NT). Decreased and increased proteins are represented in blue and red, respectively (Two-tailed t-test p-value: 0.05; log2 fold-change cutoffs of + 0.5). b) Biological pathway analysis of dysregulated proteins of control and ALS astrocytes at 14 days treated with tumor necrosis factor-alpha (TNF) (Enrichment p-value < 0.05). Pathways derived from decreased and increased proteins are represented in blue and red, respectively. Lysosomal and cytokine signaling are the most enriched altered pathways in the control condition. Membrane trafficking and immune system perturbations are the main alterations detected in ALS astrocytes after TNF treatment. c) Hierarchical clustering analysis of altered protein abundances of control and ALS astrocytes (One-way ANOVA p-value: 0.05; Tukey ’s HSD FDR: 0.05) using Pearson correlation distance. The abundance of the protein groups decreased and increased are represented in blue and red, respectively. d) Charts of pathway analysis derived from protein clusters linked to the TNF response (Enrichment p-value < 0.05). e) ALS mapping based on KEGG mapper color analysis. Protein abundances of ALS astrocytes at 14 days after TNF treatment were considered in this representation (Two-tailed t-test p-value: 0.05; log2 fold-change cutoffs of + 0.5). The ALS pathway map shows no significant changes in ALS astrocytes treated to TNF compared to ALS astrocytes not treated at 14 days. f) Immune mapping based on KEGG mapper color and Reactome analysis. Protein abundances of ALS astrocytes at 14 days after TNF treatment were considered in this representation (Two-tailed t-test p-value: 0.05; log2 fold-change cutoffs of + 0.5). Protein mapping highlights the increase in antigen presentation and non-canonical NF-kB pathway.

Furthermore, we compared the proteome profile of ALS and control astrocytes using hierarchical clustering (One-way ANOVA p-value: 0.05; Tukey ’s HSD FDR: 0.05). The analysis revealed a high similarity in the protein clusters of both TNFα-treated and non-treated ALS astrocytes (Figure 2c). Similar results were observed for control astrocytes (Figure 2c). In total, we identified 7 different protein clusters. Clusters 2 and 7 represent the protein changes in ALS and control astrocytes after TNFα treatment. In Cluster 2, we found alterations in the phagosome, cell adhesion, and amino acid metabolism, while Cluster 7 encompasses proteins associated with immune activation, such as antigen presentation and cytokine signalling (Figure 2d). In addition, we noticed two common clusters (Clusters 5 and 6) associated with ALS astrocytes regardless of treatment, as well as control astrocytes after TNFα treatment (Figure 2d). Cluster 5 exhibited downregulation of proteins involved in translation and RNA processing. Cluster 6 comprised proteins related to the immune response (Figure 2d), integrins and FGF signalling. Additionally, ALS and innate immune system mapping did not indicate other protein changes between TNFα-treated and non-treated ALS astrocytes besides cytoskeletal and endosomal perturbations and the increase in proteins related to antigen presentation and apoptosis (Figure e and f).

The relative quantification considering the protein ratios of ALS vs. control astrocytes, both treated with TNFα, revealed that ALS astrocytes exhibited an exacerbated activation of the NF-κB and cytokine signalling pathways, with a marked reduction in the metabolism of RNA and proteins (Figure S1a). The mapping of ALS signalling showed that ALS astrocytes tended to decrease mitochondrial and proteasome proteins while increasing autophagy (i.e., ATG13) and nuclear transport compared to the control (i.e., GLE1, NCP) (Figure S1b). A detailed analysis of the immune mapping disclosed that ALS astrocytes induced the preferential activation of NF-κB through the non-canonical pathway, increasing the expression of MHC-I molecules involved in immune surveillance and inflammation, as well as the production of chemoattractant cytokines such as IL-16 (Figure S1c). In summary, despite the common inflammatory responses triggered by TNFα, ALS astrocytes display a different reactivity profile compared to TNFα-treated WT astrocytes.

Collectively, these data show that the basal alterations of the innate immune system and mitochondrial dysfunction observed in TNFα-treated SOD1^A4V^ astrocytes did not occur in TNFα-treated control astrocytes.

### FGF4 reverses ALS astrocyte dysfunction and reactivity

Since FGF signalling can modulate various aspects of astrocyte physiology, such as reactivity and excitotoxicity [31], we evaluated the potential effect of FGFs (FGF2, FGF4, FGF16, and FGF18)[22] in reversing the altered biological pathways in ALS astrocytes. To induce FGF overexpression, we transduced astrocyte cultures with AAV particles encoding candidate FGFs and compared them to similar transduction with a control AAV vector expressing green fluorescent protein (GFP) (Figure 3a-c). We used proteomics to detect changes in protein profiles 14 days after astrocyte transduction (Table S2). First, we confirmed the detectability of FGF expression in the intracellular and released fractions using proteomics and ELISA assays (Figure 2c and Figure S2). We separately investigated the effect of FGF transductions on control and ALS conditions. Overall, we observed no significant changes in protein expression between non-treated (NT) and AAV-GFP-treated (GFP) astrocytes (Figure 3d and e), ruling out possible effects of GFP on the astrocytes proteome. Hierarchical clustering analysis (One-way ANOVA p-value: 0.05; Tukey ’s HSD FDR: 0.05) of control astrocytes showed two clusters. Cluster 1 and Cluster 2 represented groups whose proteins levels were decreased and increased, respectively (Figure 3d). Cluster 1 is mainly associated with cell adhesion, extracellular matrix interaction, and RNA metabolism, while Cluster 2 is involved in vesicle transport, cytoskeleton regulation, proteasome, and amino acid metabolism (Figure 3d). Similarly, two main clusters of proteins were found dysregulated in ALS astrocytes (Figure 3e). Cluster 1 consists of down-regulated proteins with biological functions related to glutathione metabolism, phagosome, spliceosome, innate immune system, and RNA metabolism. Cluster 2 includes increased protein groups mainly related to cytoskeleton regulation, vesicle transport, extracellular matrix interaction, and phosphatidylinositol signalling. We found that FGF4 triggered the most profound alterations compared to the other FGFs in ALS astrocytes. In contrast, control astrocytes displayed a more homogeneous protein profile in response to all FGFs. Specifically, in SOD1^A4V^ astrocytes, the FGFs transduction had an important effect on the immune system pathway. We located the specific protein perturbations generated by the FGF4 transduction in SOD1^A4V^ astrocytes in the ALS and immune system proteome maps (Figure 3h and i). ALS and immune system mapping expose that FGF4 in ALS astrocytes impact proteins associated with mitochondrial function, cytoskeleton, and cGAS-STING signalling axis regulation.

**Figure 3.**
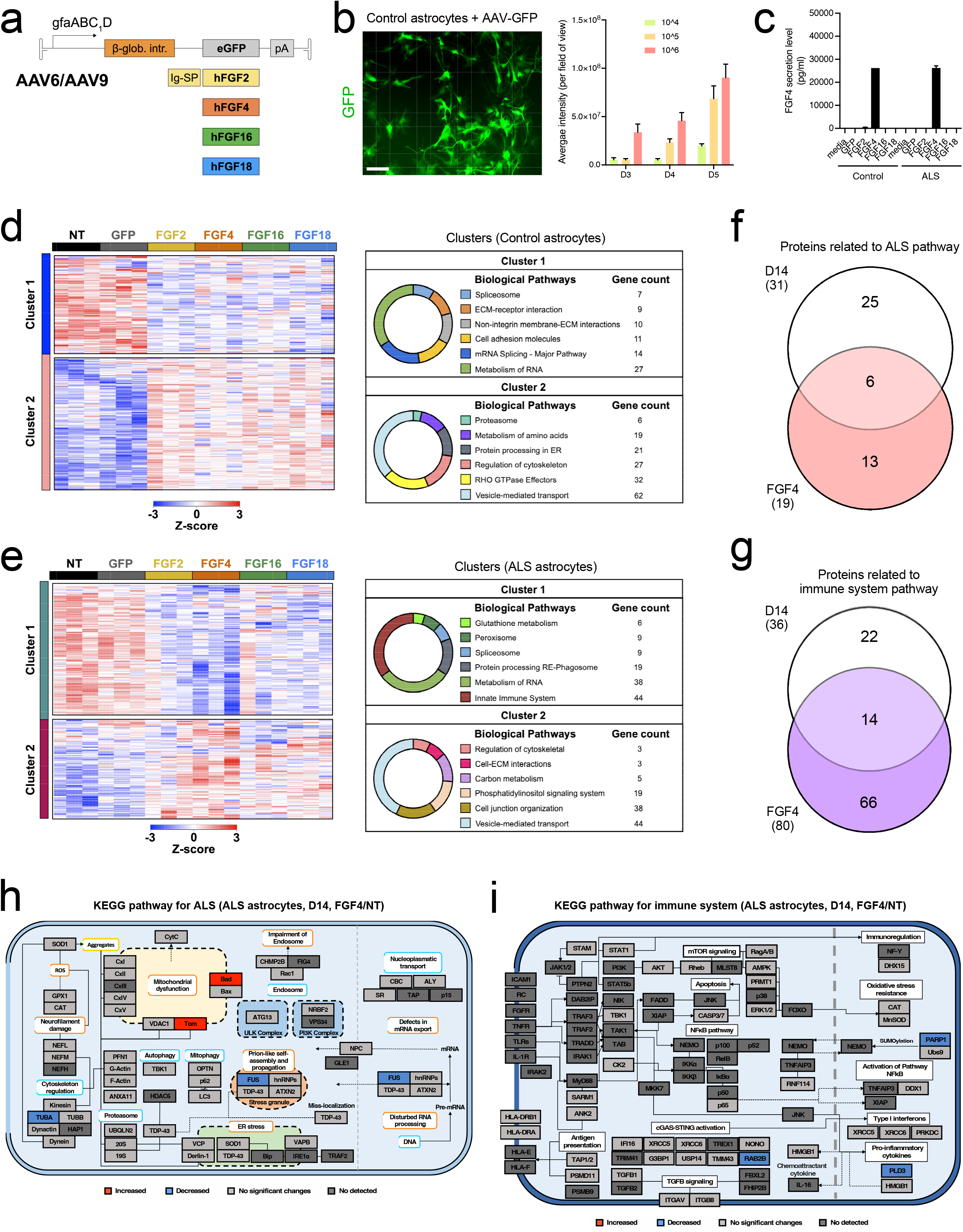
**a)** Schematic representation of AAV vectors. **b)** Astrocytes transduced with control AAV vector express GFP. Scale bar, 100 μm. The bar diagram shows levels of natural GFP in control astrocytes at 3, 4, and 5 days post-transduction with different concentrations of viruses. **c)** Bar diagram shows the quantity of FGF4 release in the medium by AAV-FGF4 transduced astrocytes, measured by ELISA (n = 2 independent experiments with triplicate wells measured for each experiment). **d** and **e)** Hierarchical clustering analysis of dysregulated proteins in control **(d)** and ALS **(e)** astrocytes treated with different FGFs (One-way ANOVA p-value: 0.05; Tukey ’s HSD FDR: 0.05) using Pearson correlation distance. The abundance of the protein groups decreased and increased are represented in blue and red, respectively. Charts of pathway analysis derived from clusters 1 and 2 (Enrichment p-value < 0.05) represent the protein groups changed after FGFs treatment. **f** and **g)** Venn diagram of dysregulated proteins related to ALS pathways and immune system pathways detected in ALS and control astrocytes on day 14 and FGF4 treated condition. **h)** ALS mapping based on KEGG mapper color analysis. Protein abundances of ALS astrocytes at 14 days after FGF4 treatment were considered in this representation (One-way ANOVA p-value: 0.05; Tukey ’s HSD FDR: 0.05). The ALS pathway map reveals a significant effect of FGF4 treatment in ALS astrocytes reversing the dysregulation of the proteins associated with the mitochondrial and proteasome function. **i)** Immune mapping based on KEGG mapper color and Reactome analysis. Protein abundances of ALS astrocytes at 14 days after FGF4 treatment were considered in this representation (One-way ANOVA p-value: 0.05; Tukey ’s HSD FDR: 0.05). Protein mapping shows the impact of FGF4 treatment on ALS astrocytes reversing the protein changes linked to the activation of the innate immune system.

Thus, we wondered if FGF4 could act on the same innate immune-related proteins previously detected as dysregulated in ALS astrocytes on D14. We found an overlap between the proteins affected by FGF4 treatment and all immune system-related proteins previously found altered in ALS astrocytes on D14 (Figure 3g). We observed the same for dysregulated proteins of the ALS pathway (Figure 3f). Then, we verified that FGF4 treatment could counteract some changes observed in ALS astrocytes on D14, returning them close to baseline levels (Figure S3).

We also assessed the responsiveness of ALS versus control astrocytes to FGF4 treatment (Figure S4 a-c). Data analysis suggests that ALS astrocytes treated with FGF4 had reduced levels of glycolysis/gluconeogenesis, protein and glutathione metabolism, and some immune-related proteins (e.g., RAB2B, XRCC5, XRCC6). However, they showed an exacerbated response in the modulation of RNA metabolism and interferon signalling (e.g., YTHDF2, SAMHD1). Proteins involved in apoptosis activation and the mitochondrial complex I were increased in SOD1^A4V^ astrocytes, while the abundance of proteins related to cytoskeleton regulation, proteasome, ER stress, and other components of the electron transport chain decreased.

Altogether the data suggest that FGF4 transduction could be a good candidate for fine-tuning the immune response of ALS astrocytes in vivo.

### FGF4 lowers ALS astrocyte reactivity in vivo but fails to protect ALS motor neurons from death

We next sought to investigate whether delivering FGF4 in vivo could influence glial reactivity and motor neuron degeneration. We injected SOD1^G93A^ mice at 60 days of age (pre-symptomatic stage) with AAV serotype 9 encoding for GFP and FGF4 intrathecally (Figure 4a). To restrict the expression of the candidate transgenes specifically to astrocytes, we used the gfaABC1D promoter [32, 33]. The spinal cord of mice from different cohorts was collected at 140 days (symptomatic stage [34]). We first confirmed successful delivery of the transgene using the GFP reporter vector. The reporter expression was limited to astrocytes in the white and grey matter of the spinal cord (Figure 4b-d).

**Figure 4.**
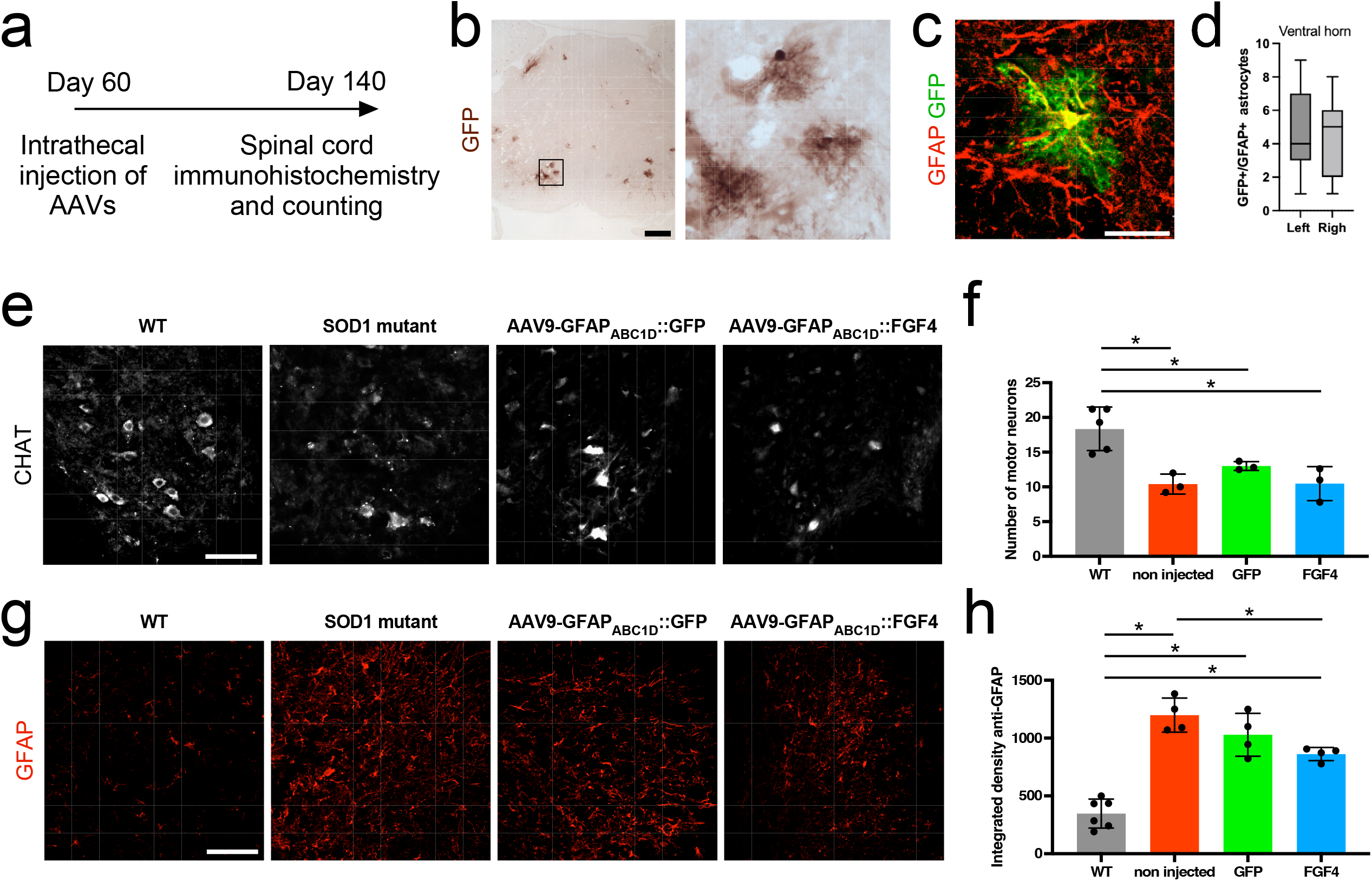
**a)** Experimental workflow: 60-day-old SOD1G93A mice were intrathecally injected with AAVs, and the spinal cord was collected at days 90 and 140 to perform immunohistochemistry. **b)** Representative image of the spinal cord stained for GFP, 30 days following intrathecal injection of AAV9-gfaABC1D-GFP. At higher magnification (right panel), GFP-positive cells show a highly complex morphology typical of astrocytes. Scale bar, 200 μm. **c)** Immunostaining for GFAP (in red) and GFP (in green) cells in the spinal cord of 90-day-old SOD1 mutant mice injected with AAV9-gfaABC1D-GFP. Scale bar, 25 μm. **d)** Box-and-whiskers plot of the number of GFP+GFAP+ astrocytes observed on each side of the ventral horn of the spinal cord. Mice were intrathecally injected with AAV9-gfaABC1D-GFP at day 60 and spinal collected 30 days later (n = 4). **e)** Representative images of the ventral horn of the spinal cord immunostained for ChAT (white) from the different experimental groups. Scale bar = 100 μm. **f)** Quantification of the number of ChAT+ motoneurons in the lumbar spinal cord of 140-day- old mice injected or not with AAV9-gfaABC1D-GFP and AAV9-gfaABC1D-FGF4 (n = 3-5). Values are mean ± SEM, *P < 0.05, one-way ANOVA with Tukey’s post hoc test. **g)** Representative immunofluorescence staining of GFAP in the spinal cord of 140-day-old wildtype, SOD1G93A, and SOD1G93A mice injected with the indicated AAVs. Scale bar = 100 μm. **h)** Bar graph of the GFAP fluorescence signal intensity in the ventral horn of the spinal cord of mice of indicated genotype, injected or not with AAV9-gfaABC1D-GFP and AAV9-gfaABC1D- FGF4. n = 3-5, values are mean ± SEM, *P < 0.05, one-way ANOVA with Tukey’s post hoc test.

Next, we assessed the survival of MNs by counting the number of choline acetyltransferase-positive neurons in the ventral horn of the lumbar SC of SOD1^G93A^ mice injected (or not) with AAV9-gfaABC1D::GFP and AAV9-gfaABC1D::FGF4 as well as age-matched WT mice (Figure 4e and f). At the symptomatic stage, we observed a significant loss of MN in non-injected SOD1^G93A^ mice and those injected with AAV9-gfaABC1D::GFP compared to WT mice. However, we did not find any neuroprotective effect of FGF4 on MNs.

We then examined the extent of astrocytic response under different experimental conditions. Glial reactivity can be quantified *in vivo* by measuring GFAP intensity. We, therefore, quantified the mean fluorescence intensity of GFAP in the lumbar SC of WT, SOD1^G93A^, and SOD1^G93A^ mice injected with different vectors (Figure 4g and h). We observed a significant increase in glial response in SOD1^G93A^ mice with AAV9-gfaABC1D::GFP vector compared to WT mice. However, local expression of FGF4 significantly reduced astrocyte reactivity in SOD1^G93A^ mice (Figure 4h).

### ALS astrocyte reactivity rescued by FGF4 is attenuated by TNFα

Work *in vivo* showed that FGF4 partially rescued astrocyte reactivity in ALS. We hypothesized that a complete rescue could not be achieved due to ongoing complex neuroinflammatory processes involving other cell types. To gain further insight, we examined how the astrocyte proteome was modulated when cells were treated with TNFα+FGF4. Volcano plots showed a consistent increase in proteins related to antigen presentation in both control and ALS astrocytes treated with TNFα+FGF4 (Figure 5a), along with a substantial increase in proteins linked to oxidative stress in the ALS astrocytes (*e.g*., SOD2). The biological pathway analysis of the control condition (Figure 5b, left panel) indicated a decline in the proteins involved in the apoptotic process, lysosome function, and keratan sulfate degradation accompanied by an increase in proteins related to the immune system, membrane trafficking, and mRNA surveillance. In contrast, the ALS astrocytes showed an underrepresentation of the membrane trafficking pathway, cytokine signalling, and ubiquitin-mediated proteolysis, while simultaneously exhibiting an increase in immune-related proteins and proteins involved in RNA and amino acid metabolism (Figure 5b, right panel). These findings suggest that in response to FGF4, astrocytes carrying SOD1^A4V^ mutant could modulate some aspects of inflammation connected to cytokine signalling. However, they are unable to completely inhibit the activation of the immune system and ALS-related biological processes.

**Figure 5.**
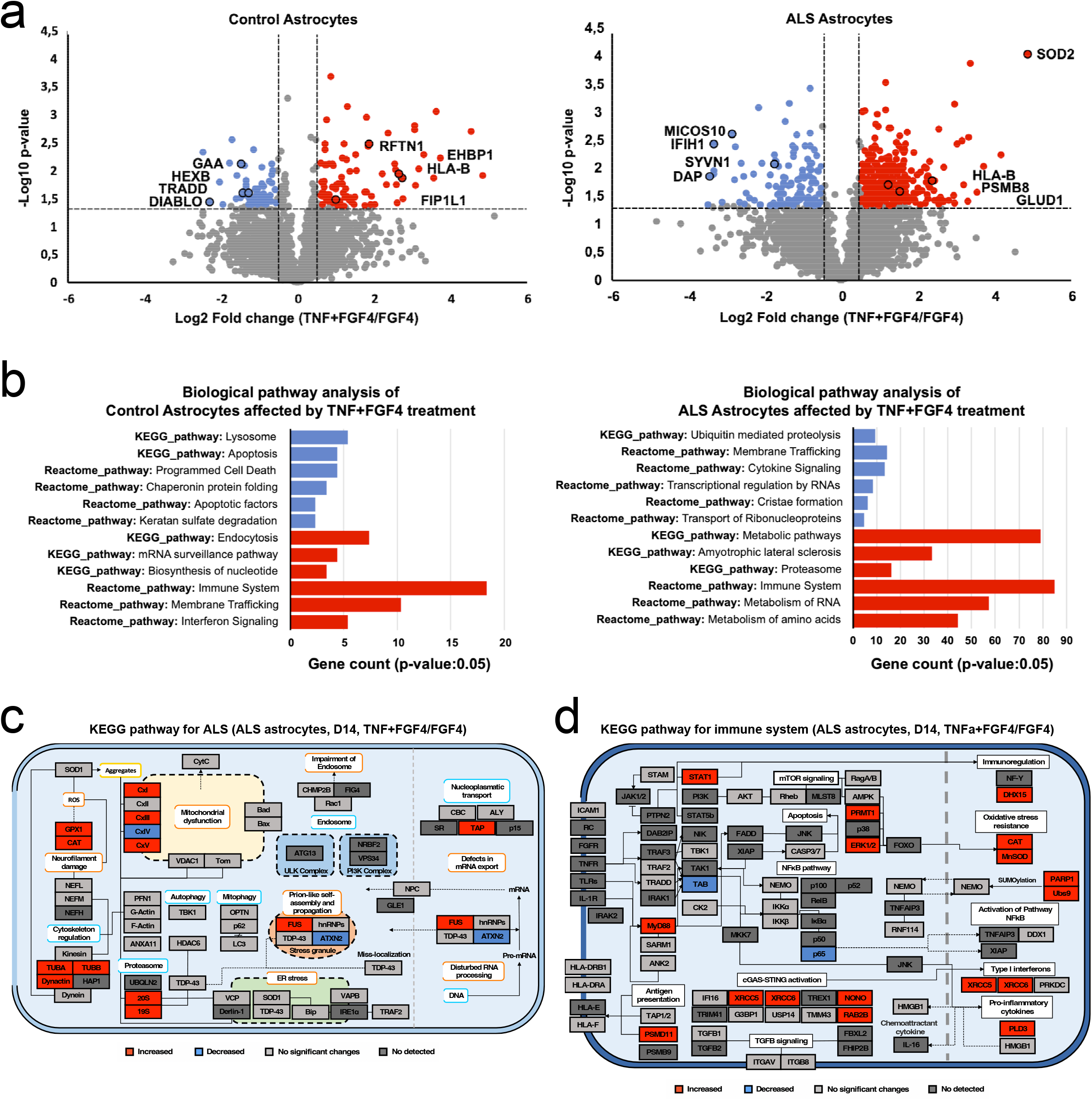
**a)** The volcano plot shows the proteome of control and ALS astrocytes at 14 days treated with TNF+FGF4 compared to FGF4 alone. Decreased and increased proteins are represented in blue and red, respectively (Two-tailed t-test p-value: 0.05; log2 fold-change cutoffs of + 0.5). **b)** Biological pathway analysis of protein changes in control and ALS astrocytes at 14 days treated with TNF+FGF4 vs. FGF4 alone (Enrichment p-value < 0.05). The decreased and increased proteins are represented in blue and red, respectively. Lysosomal and immune system signaling are the most enriched altered pathways in the control condition. In contrast, membrane trafficking and the immune system remain the main dysregulated pathways in ALS astrocytes. **c)** ALS mapping based on KEGG mapper color analysis. Protein abundances of ALS astrocytes at 14 days treated with TNF+FGF4 versus FGF4 were considered in this representation. The p-value of the two-tailed t-test is 0.05 with log2 fold-change cutoffs of + 0.5. The data show that under a pro-inflammatory environment, the treatment with FGF4 fails in reverting the mitochondrial and proteasome dysregulated proteins. **d)** Immune mapping based on KEGG mapper color and Reactome analysis. Protein abundances of ALS astrocytes at 14 days treated with TNF+FGF4 vs. FGF4 were considered in this representation (Two-tailed t-test p-value: 0.05; log2 fold-change cutoffs of + 0.5). Protein mapping confirms that FGF4 treatment in the presence of TNF partially reduces the protein changes linked to immune system activation.

We proceeded to determine the specific protein changes connected to TNFα+FGF4 treatment in the ALS astrocytes, locating these perturbations in the ALS and immune system maps (Figure 5c and d). SOD1^A4V^ astrocytes increment some components of the mitochondrial respiratory complex, proteasome, and antioxidant enzymes such as catalase and glutathione peroxidase 1. We also observed an increase of protein levels involved in NEMO SUMOylation (i.e., PARP1, UBS9), innate immune response regulation (e.g., PRMT1), activation of the cGAS-STING pathway (e.g., XRCC5, XRCC6, NONO, RAB2B), and Toll-like receptor (TLR) signalling (e.g., MYD88, PLD3). Additionally, the effect of TNFα+FGF4 treatment on ALS astrocytes stimulated the increment of proteins related to the regulation of TGFB signalling (e.g., SDCBP, AIMP1, HSP90AB1) and important secreted factors such as Macrophage migration inhibitory factor (MIF).

Finally, we compared the responsiveness of ALS versus control astrocytes to TNFα+FGF4 treatment (Figure S5a-c). Data analysis showed that ALS astrocytes have an exacerbated response incrementing the innate immune-related proteins, apoptosis regulation, and lysosomal and proteasomal function. However, there was a decrease in membrane traffic, mRNA surveillance, and cytokine pathways. SOD1 astrocytes also show an inclination towards increased levels of proteins associated with mitochondrial dysfunction, oxidative stress, proteasome function, and mitophagy (Figure S5b). Additionally, we observed an elevation in proteins that act with the mTOR signalling pathway and apoptotic proteins, such as caspases 3 and 7. Interestingly, our immune mapping revealed that ALS astrocytes treated with FGF4 under inflammatory condition were less susceptible to activating NF-κB, TGFβ signalling (e.g., TGFβ2). We also noted a decreased in proteins involved in cGAMP transport, such as LRRC8D, and FGF signalling (i.e., ITGB3), were also decreased (Table S2). ALS astrocytes also preserved a higher level of proteins linked to the innate immune response regulation via cGAS-STING (e.g., NONO, RAB2B) NF-κB pathway regulators (e.g., DDX1, RNF114, TNFAIP3), TLR signalling (e.g., MYD88, PLD3) and complement regulation including complement factor I (Table S2). Furthermore, we observed alterations in the levels of proteins involved in regulating inflammation and cell-intrinsic initiation of autoimmunity (e.g., TREX1) were observed.

Collectively, these data show that ALS astrocytes responded differently to TNFα insult and FGF4 transduction compared to control astrocytes. FGF4 did not completely block the activation of the innate immune system via the cGAS-STING pathway. Under pro-inflammatory conditions, FGF4 treatment modulated the signalling of certain inflammatory pathways, such as NF-κB, and antigen presentation. However, FGF4 could not prevent the increase of mitochondrial complex, apoptotic, and proteasome machinery pathways mediated by TNFα insult, nor the imbalance in TGFβ and TLR signalling and complement. Despite these limitations in neutralizing molecular changes driven by the TNFα treatment, FGF4 seemed promising in modulating essential aspects of the neuroinflammatory process *in vitro*, such as the activation of NFκβ, and MHC-I presentation.

## Discussion

Accumulated evidence suggests that neuroinflammation is one of the main drivers of MNs degeneration in ALS [35]. Astrocytes display an inflammatory profile in the pre-symptomatic phase of the disease, making them as key players in the non-cell autonomous mechanisms underlying ALS progression. Here, we demonstrate that mutant SOD1^A4V^ astrocytes exhibit a higher basal expression of proteins associated with the activation of the innate immune system as well as mitochondrial dysfunction. The increase in mitochondrial complexes is linked to the production of oxygen-reactive species (ROSs) [36]. An excess of ROSs is associated with the opening of the mitochondrial permeability transition pore (mPTP) [37] and the cytosolic escape of mitochondrial DNA (mtDNA), triggering the activation of innate immunity [38]. We observed in our data the increase of one of the main proteins involved in this phenomenon, VDAC1 [29, 39]. Concomitantly, we observed an increase in cytosolic DNA sensors such as XRCC5/XRCC6 [40], proteins necessary for the efficient activation of cGAS (i.e., G3BP1) [41], and modulators of the STING signaling at the ER membrane (i.e., TMEM43) [42]. All this evidence strongly suggests that this pathway is an important early pathogenic feature of mutant SOD1^A4V^ astrocytes. In this context, our results align with recent studies that have reported the mitochondrial release of mtDNA and the subsequent activation of the cGAS-STING-pathway in TDP-43 [43], and SOD1 mutated animal models [44]. We corroborated that the TNFα supply evoked an exacerbation of the immune pathways related to antigen presentation and NF-κB activation. However, TNFα failed to emulate in control astrocytes the perturbation of proteins associated with the innate immune response and the mitochondrial dysfunction detected in ALS astrocytes, pointing out that these are essential molecular signatures behind the ALS astrocyte phenotype.

Some FGF family members have been previously associated with the improvement of astrocyte functions, such as the increase of GLT1 expression [22, 23] and immunomodulation in different inflammatory diseases [45]. When SOD1^A4V^ astrocytes were transduced with FGFs, it primarily affects proteins related to the immune system. We have demonstrated that ALS astrocytes have a stronger response to FGF4 transduction. FGF was downregulated in the gene expression profile of ALS SC grey matter post-mortem tissues [46]. The biological function of FGF4 is linked to growth [47], survival [48], angiogenesis [49], metabolism [50], and apoptosis regulation [51]. Growing evidence suggests that FGF4 intervenes in the modulation of the immune system in early life stages [52]. We found that FGF4 counteracts some changes in the protein profile of SOD1^A4V^ astrocytes linked to ALS and the innate immune system pathways. However, our data showed that FGF4 reduces astrocyte reactivity without impacting the survival of MNs *in vivo*.

We examined the molecular changes driven by FGF4 transduction in ALS and control astrocytes cultured under a pro-inflammatory environment. The analysis of the astrocytes’ proteomes revealed that ALS astrocytes maintained elevated levels of proteins related to the cGAS-STING pathway, such as XRCC5/XRCC6, and scaffold elements for IRF3 phosphorylation (e.g., NONO, RAB2B) [53, 54]. Previous studies have shown that the complete blocking of the cGAS-STING pathway is necessary to ameliorate ALS progression and survival *in vivo* [43, 44]. Additionally, SOD1^A4V^ astrocytes exhibited a significant increase in the mTORC signalling pathway, crucial for metabolic remodelling during the immune response [55, 56], and the increment in downstream members of the TLR and interleukin-1 receptor families such as MYD88 [57]. Similarly, ALS astrocytes TNFα+FGF4 preserved high levels of MIF, a pro-inflammatory cytokine related to chronic neuroinflammation and neurodegeneration [58, 59]. We also detected the decline of proteins such as the LRRC8D subunit intimately associated with increased cGAMP transport to neighbouring cells [60], and reduced levels of the secreted complement factor I, a regulator of complement activation [61, 62]. Our data further showed that ALS astrocytes treated with TNFα+FGF4 had perturbations in the TGFβ signalling pathway, which has previously been linked to ALS progression [63]. The reduction of TGFβ-2 has been associated with impairments in the synaptic function [64], and it has been shown that a high dose of TGFβ-2 transitory improves motor performance in SOD1 mutant mice [65]. Our data suggest that ALS astrocytes TNFα+FGF4 reduce the activation of NF-κβ and antigen presentation but fail to protect MNs *in vivo* due to a complex interplay of immune system signalling.

In conclusion, this study showed the potential of the patient-derived iPSC models to capture the cell autonomous contribution of astrocytes in the pathogenesis of ALS and testing new therapeutic approaches to fine-tune the responsiveness of astrocytes to modulate the neuroinflammation in ALS.

## Experimental procedures

All materials were purchased from Thermo Fisher Scientific, unless specified.

### Human iPSC Lines

Human IPSC lines (Table S3) were obtained from RUCDR Infinite Biologics, where the iPSCs were generated by reprogramming of ALS patient and healthy control fibroblasts with Sendai or retroviral vectors and characterized for expression of common pluripotency markers and ability to differentiate into the three germ layers. All obtained iPSC lines had normal karyotype. The iPSCs were further expanded and bio-banked. The generation of iPSC line CSC-37N was previously published [66].

### Generation of spinal cord astrocytes

The iPSCs lines were differentiated to SC astrocytes using the modified protocol originally developed by Roybon et al. [23]. Control line FA11 underwent two independent differentiations. Human iPSC colonies were harvested and plated to ultra-low attachment flasks (Corning) in WiCell media supplemented with 20 μM of ROCK inhibitor Y27632 (Selleck Chemicals, Munich, Germany) and 20 ng/mL of FGF2 (Peprotech). Next day, the media was replaced by WiCell media supplemented with 0.1 μM LDN193189 (Stemgent), 10 μM SB431542 (Sigma-Aldrich), and 1 μM retinoic acid (RA; Sigma-Aldrich). On D4, 50% of WiCell media were replaced with medium composed of advanced DMEM/F12, 2 mM L-glutamine, 1% NEAA (v/v), 2% B27 Supplement without vitamin A (v/v), 1% penicillin-streptomycin (v/v), heparin, plus LDN193189, SB431542 and RA. On D6 1 μM RA, 1 μM SAG (Selleck Chemicals), 10ng/ml BDNF (Peprotech) and 0,4 μg/ml ascorbic acid (Sigma-Aldrich) were added to the mix of 50% WiCell/ 50% advanced DMEM/F12 media, at D8 and D10 to advanced DMEM/F12 and at D12- D30 to the mix of 50% advanced DMEM/F12/50%Neurobasal media with medium change every other day. In addition, on D12-30, 20 ng/ml GDNF (Peprotech) was added to the media. From D30 onwards, the free-floating spheres were grown in media composed of advanced DMEM/F12, 2 mM L-glutamine, 1% NEAA (v/v), 2% B27 supplement without vitamin A (v/v), 1% penicillin-streptomycin (v/v) and heparin in the presence of 20 ng/ml FGF2 (Peprotech) and 20 ng/ml EGF (Peprotech). On D60, free-floating spheres were collected, washed, dissociated using accutase (Invitrogen) and single cells seeded and grown as adherent cultures in flasks coated with 100 μg/ml Poly-L-ornithine (Sigma-Aldrich) and 5 μg/ml mouse laminin in media composed of advanced DMEM/F12, 2 mM L-glutamine, 1% NEAA (v/v), 2% B27 supplement without vitamin A (v/v), 1% penicillin-streptomycin (v/v), heparin and 20 ng/ml CNTF (Peprotech). From day 85, astrocytes were cultured in neural differentiation medium composed of neurobasal media, 2 mM L-glutamine, 1% NEAA (v/v), 1% N2 Supplement (v/v), 1% penicillin-streptomycin (v/v), and 20 ng/ml CNTF.

### Viral production and transduction

FGF-encoding constructs were generated by replacing, using standard cloning procedures, the GFP-coding sequence with cDNAs encoding the human forms of FGF4 (NM_002007.4), FGF16 (NM_003868.3) and FGF18 (NM_003862.3) in the pAAV-gfaABC1D-β-globin intron-GFP vector [67]. To enhance FGF2 secretion, the vector was adapted for N-terminal fusion of the protein to the immunoglobulin signal peptide, as previously described [68]. For transgene expression in astrocytes *in vivo*, each of these constructs was packaged into serotype-9 AAV particles. To transduce iPSC-derived astrocytes, the same constructs were packaged into serotype-6 AAV. Briefly, for packaging into AAV vectors, HEKExpress® cells maintained in suspension were transiently transfected with pAAV and either the pDP6 and pDP9 helper plasmid. To recover released AAV particles, the cell supernatant was collected at day 3 and day 7 after transfection. At day 7, the packaging cells were lysed using repeated freeze-thawing. For purification and concentration, AAV particles were captured by affinity chromatography using POROS™ CaptureSelect™ resins (AAV9 and AAVX, Thermo Fisher Scientific, MA). AAV suspension was concentrated using centrifugal devices (Amicon® Ultra-15 100 kDa centrifugal filter devices, Merck) and resuspended in PBS with 0.001% Pluronic-F68 before storage. The vectors were tittered by dPCR (QIAcuity® digital PCR system, QIAGEN) using an amplicon located in the β- globin intron. Overall, the method of AAV vector production was adapted from the previously described protocol [69]. To overexpress FGFs in human astrocytes, astrocytes aged days 98 – 107 were seeded on the plate coated with 100 μg/ml poly-L-ornithine and 5 μg/ml mouse laminin. After 3 hours, the astrocytes were transduced with AAV6-GFAP-FGF2, -FGF4, -FGF16 or - FGF18 at 2x10(5) VG/cell. At D5 or D14 post-transduction, the cells were collected, washed with PBS and frozen at -80C for further analysis.

### TNFα treatment

Astrocytes aged days 98 – 107 were seeded in 24 well plate coated with 100 μg/ml P/O and 5 μg/ml mouse laminin at the density of 120,000 cells/well. 48h after, the cells were treated with either 100 ng/ml TNFα, or 100 ng/ml TNFα and AAV vector expressing FGF4 for 14 days. After that, the cells were washed with PBS and frozen at −80°C until analyzed.

### Immunocytochemistry

For immunostaining, cultures were fixed using 4% paraformaldehyde for 20 min at room temperature and nonspecific binding was blocked with 10% normal donkey serum and 0.1% Triton X-100 in PBS for 1hour. The cells were then incubated with primary antibody to GFAP (1:1000, #Z0334, DAKO), S100b (1:500, #S2532, Sigma-Aldrich), ezrin (1:500, #3145S, Cell Signaling), GS (1:500, #MAB-302, Millipore) and ID3 (1:500, #9837S, Cell Signaling) overnight at 4°C. Next day the cells were washed and incubated with Alexa Fluor 488 nm- (1:400; #A- 21202, Molecular Probes) or 555 nm-conjugated secondary antibodies (1:400; #A-31570, Thermo Fisher Scientific) for 1 hr. In addition, the cells were counterstained with DAPI (1:10,000, Life Technologies). All images were obtained using an inverted epifluorescence microscope (LRI – Olympus IX-73).

### Enzyme-linked immunosorbent assay (ELISA)

The FGFs secretion in culture media was measured by ELISA D5 post-transduction of the cultures according to the manufacturer’s instructions; FGF2, FGF4, FGF16 (all three from Thermo Fisher Scientific), FGF18 (LSBio, Seattle, WA, USA). Absorbance was measured at 450 nm.

### Protein extraction and sample digestion

Proteins from astrocytes at days 5 and 14 were extracted using a lysis buffer of 25mM DTT, 10 w/v% SDS in 100mM triethylammonium bicarbonate (TEAB). The cells were sonicated using 40 cycles of 15s on/off at 4°C in the Bioruptor plus (model UCD-300, Diagenode). Further, the samples were boiled at 99°C for 5 minutes and centrifuged at 20,000 g for 15 min at 18°C. The supernatants were collected to determine the protein amounts (Pierce 660nm Protein Assay with Ionic Detergent Compatibility Reagent). Proteins were preserved at -80°C until further use. The proteins were digested using S-Trap™ 96-well plate following the instructions of the manufacturer (ProtiFi S-Trap^TM^ 96-well plate digestion protocol). Briefly, samples were alkylated with 50 mM IAA for 30 min in the dark at room temperature, followed by tryptic digestion in 50 mM TEAB (enzyme: substrate,1:50) at 37°C overnight. The peptides were eluted in three steps, first with 80 µL of 50 mM TEAB, then with 80 µL of 0.2% formic acid (FA), and finally with 80 µL of 50% acetonitrile (ACN) containing 0.2% FA. Peptides were dried in a speed-vac and resuspended in 0.1% trifluoroacetic acid (TFA)/2% ACN for peptide concentration measurement (Pierce Quantitative Colorimetric Peptide Assay).

### nanoLC-MS/MS analysis and Database search

The nLC-MS/MS analysis was performed on a QExactive HF-X mass spectrometer coupled to a Dionex Ultimate 3000 RSLC nano UPLC system (Thermo Scientific), with an EASY-Spray ion source. 500 ng of peptides was analyzed from each sample using high-resolution data independent acquisition (HR-DIA). All samples were loaded onto an Acclaim PepMap 100 C18 (75 µm × 2 cm, 3 µm, 100 Å, nanoViper) trap column and separated on an Acclaim PepMap RSLC C18 column (75 µm × 50 cm, 2 µm, 100 Å) (Thermo Scientific) using a flow rate of 300 nL/min, a column temperature of 60°C. A 110 min gradient was applied for separation, using solvents A (0.1% formic acid) and B (0.1% formic acid in 80% ACN), increasing solvent B from 2 to 45% in 95 min and to 95% in the next 15 min, continuing for another 5 min. In HR-DIA analysis, a complete acquisition cycle consisted of 3 MS1 full scans, each followed by 18 MS2 DIA scans with variable isolation windows. Full MS1 scans were acquired using a mass range of 375-1455 m/z, with a resolution of 120,000 (at 200 m/z), target AGC value of 3x10^6^ and maximum injection time of 50 ms. MS2 scans were acquired with a resolution of 30,000 (at 200 m/z), target AGC value of 1x10^6^, automatic maximum injection time, fixed first mass of 200m/z, NCE of 28. The variable isolation windows were 13, 16, 26 and 61 m/z with 27, 13, 8, 6 loop counts, respectively. Direct DIA analysis of the raw files was performed in Spectronaut v14 (Biognosys) against the reviewed Uniprot *Homo sapiens* sequence database (May 2022, 42363 entries) using BGS factory default settings. The identifications were filtered by a Q value of <0.01 (equals a FDR of 1% on peptide level). Quantification was done on MS1 level, using the top3 method.

### AAV injection in mutant SOD1 mice model of ALS

B6.Cg-Tg(SOD1*G93A)1Gur/J (*SOD1^G93A^*) mice were purchased from the Jackson Laboratory and maintained on a C57BL/6 background. Each vector was injected at a titer of 3.10^14^ vg/ml. A volume of 10 µl of viral vectors mixed with 5 µl of lidocaine (10 mg/ml) was slowly injected between the L5 and L6 vertebraes. A total of 4 mice per condition were injected. Appropriate injection into the intrathecal space was confirmed by the animal’s tail movement and by the transiently anesthetized lower limbs.

### Immunohistochemistry

Mice were euthanized at 140 days of age by intra-peritoneal injection of ketamine and xylazine. The mice were then transcardially perfused with PBS and the SC was incubated in 4% paraformaldehyde in PBS for 45 min at +4°C. After incubation, the SC was transferred to a 30% sucrose solution and imbedded in Tissue-Tek OCT compound (Sakura Finetek). For immunofluorescence labeling, cryosections of the lumbar SC (16 μm) were collected onto Superfrost Plus glass slides. The sections were then incubated in PBS containing 0.05 % Tween-20, 0.3% Triton-X100 and 20% heat-inactivated donkey serum (blocking solution) for 1 h at room temperature (RT). Next, sections were incubated for 24 h at +4°C with primary antibodies (goat anti-Chat (1:200, Merck, AB144P), mouse anti-GFAP (1:300, Merck, MAB360), and chicken anti-GFP (1:700, Abcam, Ab13970) diluted in the blocking solution. Sections were washed with PBS and incubated with fluorochrome-conjugated secondary antibodies diluted in the blocking solution for 1 h at RT. Sections were washed with PBS and mounted onto glass slides using NeoMount Fluo (NeoBiotech) solution. For chromogenic immunostaining, cryosections were incubated in Tris-buffered saline (TBS) containing 0.3 % Triton-X100 and 1.5 % donkey serum (blocking solution) for 1 h at RT. The samples were then incubated overnight at +4°C with the primary antibody (rabbit anti-GFP, 1:600, Chromotek) diluted in the blocking solution. Sections were washed with TBS and incubated with biotinylated secondary antibody (1:100; Vector labs) diluted in blocking solution for 1 h at RT. Following several washes with 0.05 % Tween-20 in TBS, the tissue was incubated with ABC complex (Vectastain ABC kit, Vector labs kit) in 0.05 % Tween-20 in PBS for 1 h. After washing with 0.05 % Tween-20 in TBS and TB, sections were incubated with the 3,3’ diaminobenzidine substrate solution according to the manufacturer’s instructions. Image acquisitions were performed using a Zeiss Axio imager microscope. For astrocytic activation, image acquisition was done on a Zeiss LSM880 Airyscan confocal microscope. The GFAP signal intensity was determined with ImageJ (National Institutes of Health).

### Statistical and biological pathway analyses

Protein abundances were normalized by median subtraction following log2 transformation. Principal component and statistical analyses were performed using the Perseus 1.6.5.0 software. To compare only two conditions, the two-tailed t-test (p-value:0.05) with a cutoff of log2 ± 0.5-fold-change was applied. One-way ANOVA (p-value:0.05) and Tukey’s HSD (FDR: 0.05) were done for multiple comparisons. The hierarchical clustering analysis was performed using the Pearson’s correlation distance. The Functional Annotation Tool DAVID Bioinformatics Resources was used to detect the relevant biological pathways, considering Reactome Gene Sets and KEGG Pathway (p-value: 0.05).

### Declarations

#### Ethical Approval and Consent to participate

All animal experiments were approved by the national ethics committee on animal experimentation and were done in compliance with the European community and national directives for the care and use of laboratory animals.

#### Consent for publication

Not applicable.

#### Availability of supporting data

The mass spectrometry proteomics data have been deposited to the ProteomeXchange Consortium via the PRIDE [70] partner repository with the dataset identifier PXD045267.

#### Competing interests

The authors declare that they have no competing interests.

#### Funding

We acknowledge funding support from the strategic research areas MultiPark at Lund University. This work was supported by The French Muscular Dystrophy Association (*AFM*- *Téléthon*), The Crafoord Foundation, The Åhlens Foundation, and The Olle Engkvist Byggmästare Foundation. E.V. was supported by The Olle Engkvist Byggmästare Foundation,

E.S. was supported by The French Muscular Dystrophy Association (*AFM*-*Téléthon*). S.M-M-A. is supported by the Foundation for Medical Research (FRM).

#### Authors’ contribution

E.V., E.S. and L.R. conceived the experiments and wrote the manuscript; all authors performed or assisted with the experiments; and provided reagents, expertise, and conducted a critical analysis and review of the manuscript. All authors provided input during the editing of the manuscript and approved its content.

## Acknowledgments.

We are particularly thankful to Target ALS (http://www.targetals.org/) and RUCDR Infinite Biologics (formerly Rutgers University Cell and DNA Repository) for providing the control and ALS patient iPSC cell lines. We gratefully acknowledge the support of BioMS (Swedish National Infrastructure for Biological Mass Spectrometry).

## List of abbreviations

Adeno-associated virus: (AAV)
Amyotrophic lateral sclerosis: (ALS)
Cyclic GMP-AMP synthase - Stimulator of interferon genes: (cGAS-STING)
Extracellular matrix: (ECM)
Familial ALS: (fALS)
Fibroblast growth factor 2: (FGF2)
Glial fibrillary acidic protein: (GFAP)
Glutamate transporter-1: (GLT1)
Glutamine synthetase: (GS)
Green fluorescent protein: (GFP)
Induced pluripotent stem cells: (iPSCs)
Inhibitor of DNA binding 3: (ID3)
Mitochondrial DNA: (mtDNA)
Mitochondrial permeability transition pore: (mPTP)
Motor neuron: (MN)
Oxygen-reactive species: (ROS)
Spinal cord: (SC)
Sporadic ALS: (sALS)
Superoxide dismutase 1: (SOD1)
S100 calcium binding protein beta: (S100b),
Tricarboxylic acid cycle/Electron transport chain: (TCA/ETC)
Tumor necrosis factor-alpha: (TNFa)
Voltage-dependent anion-selective channel 1: (VDAC1)
Wildtype: (WT)

## Notes

### Competing Interest Statement

The authors have declared no competing interest.

