## supporting figure legends for "TNFα hinders FGF4 efficacy to mitigate ALS astrocyte dysfunction and cGAS-STING pathway-induced innate immune reactivity"

***Figure S1***

**a)** Volcano plots of ALS vs. control astrocytes at 14 days treated with TNFa (left panel). Decreased and increased proteins are represented in blue and red, respectively (Two-tailed t-test p-value: 0.05; log2 fold-change cutoffs of + 0.5). Biological pathway analysis of astrocyte dysregulated proteins treated with TNFa (Right panel. Enrichment p-value < 0.05). Pathways derived from decreased and increased proteins are represented in blue and red, respectively.

**b)** ALS mapping based on KEGG mapper color analysis. Protein abundances of ALS vs. control astrocytes at 14 days treated with TNFa were considered in this representation (Two-tailed t-test p-value: 0.05; log2 fold-change cutoffs of + 0.5).

**c)** Immune mapping based on KEGG mapper color and Reactome analysis. Protein abundances ALS vs. control astrocytes at 14 days treated TNFa were considered in this representation (Two-tailed t-test p-value: 0.05; log2 fold-change cutoffs of + 0.5).

***Figure S2***

**a)** Relative quantification of FGFs using proteomics as a readout. Protein intensities were normalized using the subtraction of the median.

***Figure S3***

**a)** Hierarchical clustering analysis of dysregulated proteins in control, ALS astrocytes NT AND ALS+FGF4 astrocytes (One-way ANOVA p-value: 0.05; Tukey´s HSD FDR: 0.05) using Pearson correlation distance. The abundance of the protein groups decreased and increased are represented in blue and red, respectively.

**b)** Charts of pathway analysis derived from clusters 3 and 5 (Enrichment p-value < 0.05) represent the protein groups reverted in ALS astrocytes after FGF4 transduction.

***Figure S4***

**a)** Volcano plots of ALS vs. control astrocytes at 14 days treated after FGF4 transduction (left panel). Decreased and increased proteins are represented in blue and red, respectively (Two-tailed t-test p-value: 0.05; log2 fold-change cutoffs of + 0.5). Biological pathway analysis of astrocyte dysregulated proteins after FGF4 transduction (Right panel. Enrichment p-value < 0.05). Pathways derived from decreased and increased proteins are represented in blue and red, respectively.

**b)** ALS mapping based on KEGG mapper color analysis. Protein abundances of ALS vs. control astrocytes at 14 days after FGF4 transduction were represented (Two-tailed t-test p-value: 0.05; log2 fold-change cutoffs of + 0.5).

**c)** Immune mapping based on KEGG mapper color and Reactome analysis. Protein abundances ALS vs. control astrocytes at 14 days after FGF4 transduction were represented (Two-tailed t-test p-value: 0.05; log2 fold-change cutoffs of + 0.5).

***Figure S5***

**a)** Volcano plots of ALS vs. control astrocytes at 14 days treated with TNFa+FGF4 (left panel). Decreased and increased proteins are represented in blue and red, respectively (Two-tailed t-test p-value: 0.05; log2 fold-change cutoffs of + 0.5). Biological pathway analysis of astrocyte dysregulated proteins treated with TNFa+FGF4 (Right panel. Enrichment p-value < 0.05). Pathways derived from decreased and increased proteins are represented in blue and red, respectively.

**b)** ALS mapping based on KEGG mapper color analysis. Protein abundances of ALS vs. control astrocytes at 14 days treated with TNFa+FGF4 were considered (Two-tailed t-test p-value: 0.05; log2 fold-change cutoffs of + 0.5).

**c)** Immune mapping based on KEGG mapper color and Reactome analysis. Protein abundances ALS vs. control astrocytes at 14 days treated TNFa+FGF4 were considered (Two-tailed t-test p-value: 0.05; log2 fold-change cutoffs of + 0.5).
