## Supplementary figures and images for "TNFα hinders FGF4 efficacy to mitigate ALS astrocyte dysfunction and cGAS-STING pathway-induced innate immune reactivity"

### figure S1

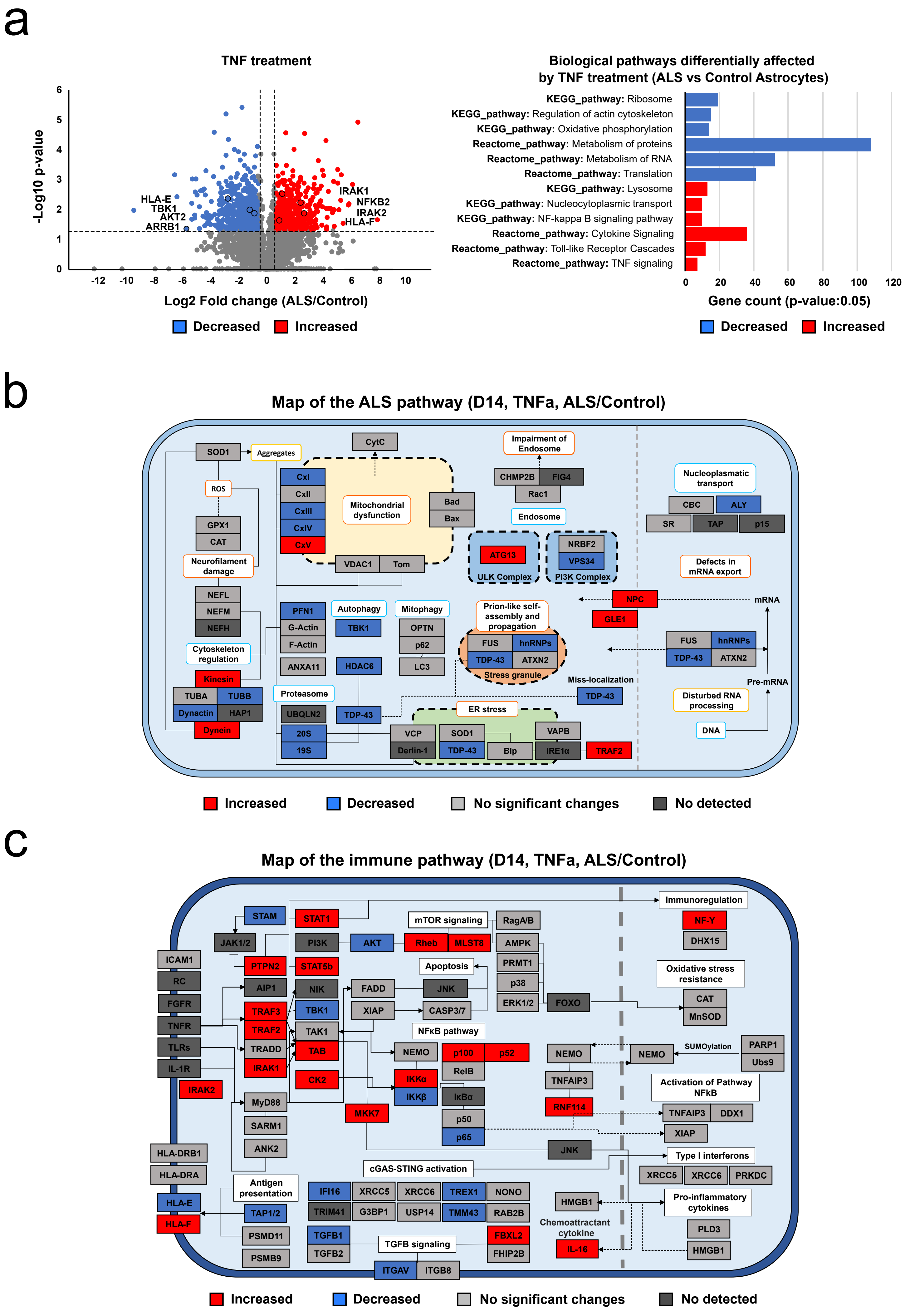

### figure S2

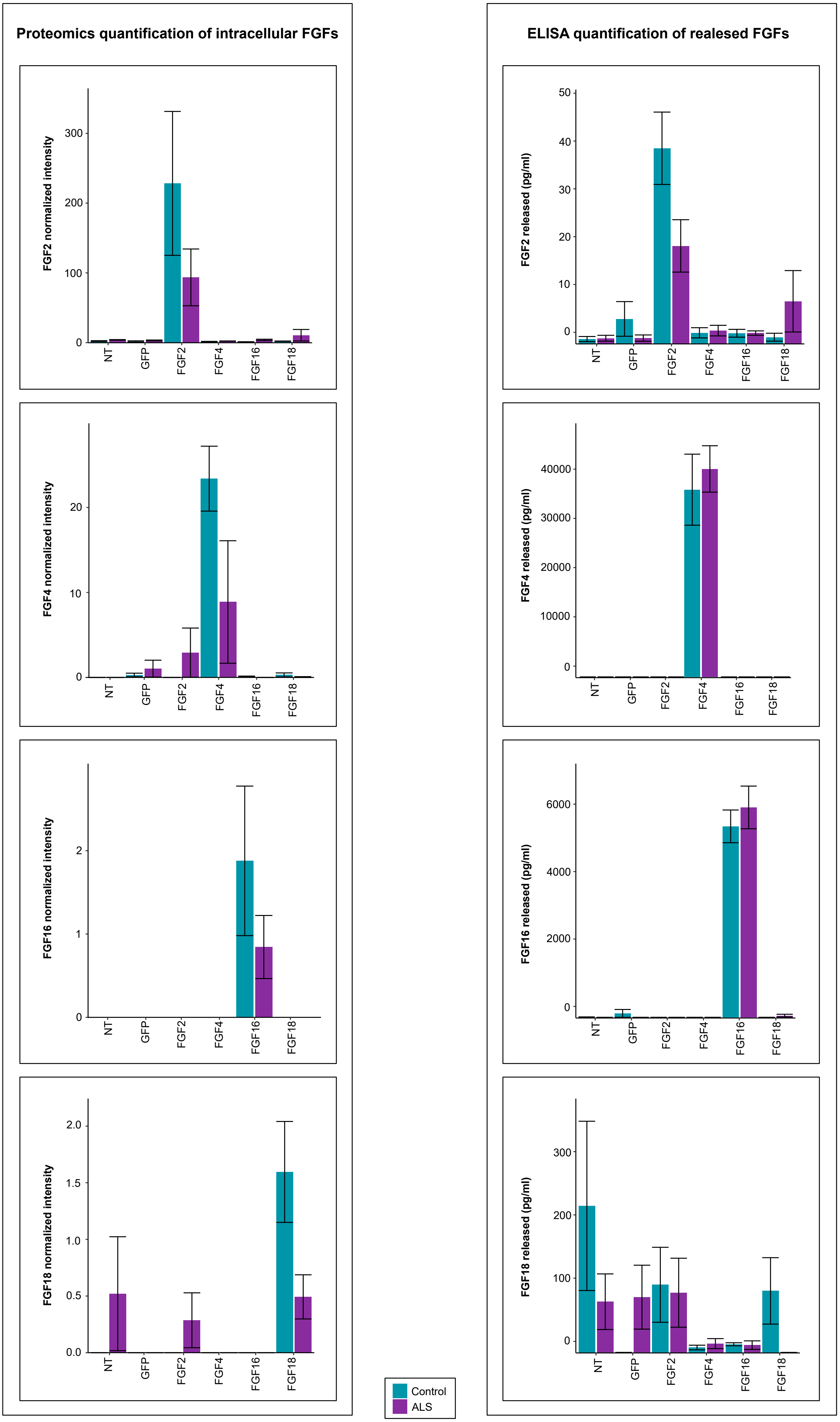

### figure S3

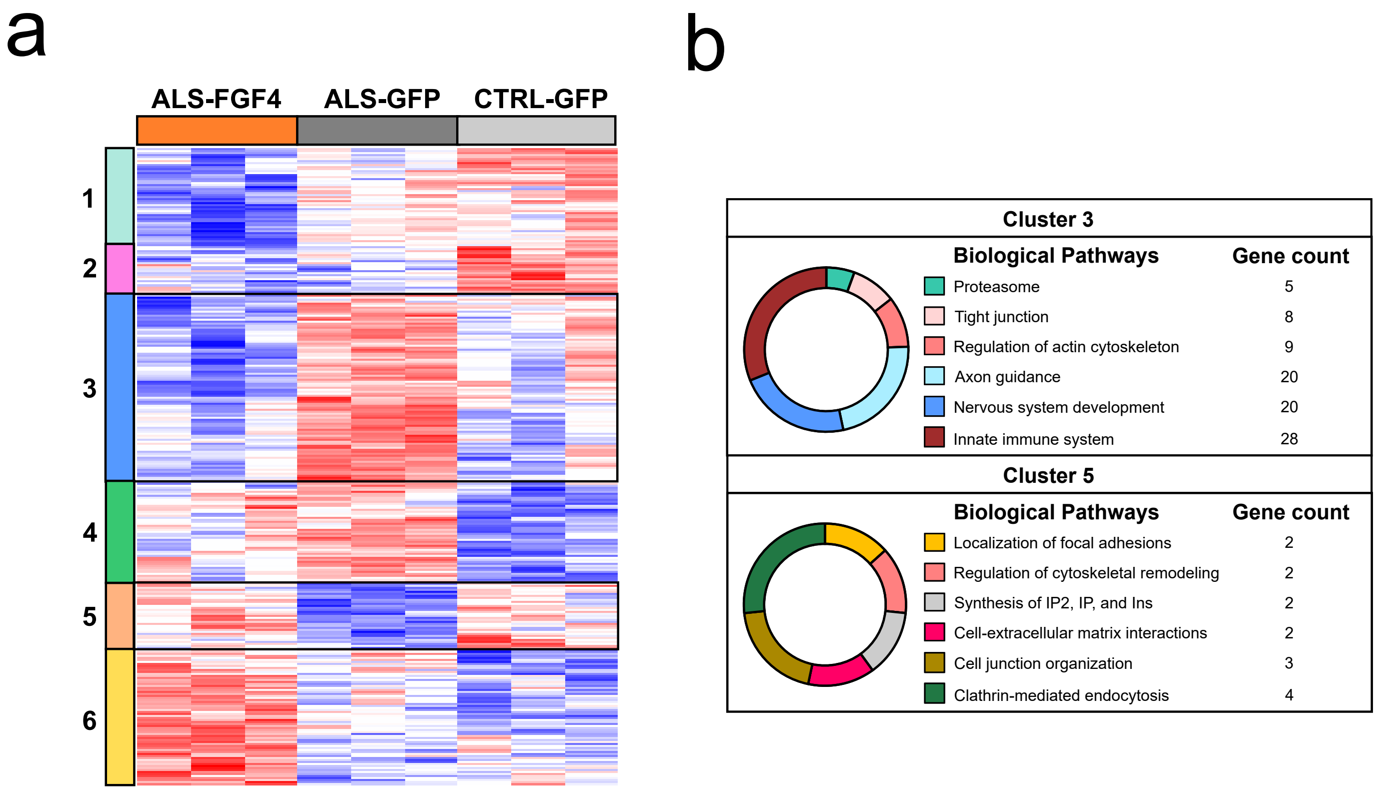

### figure S4

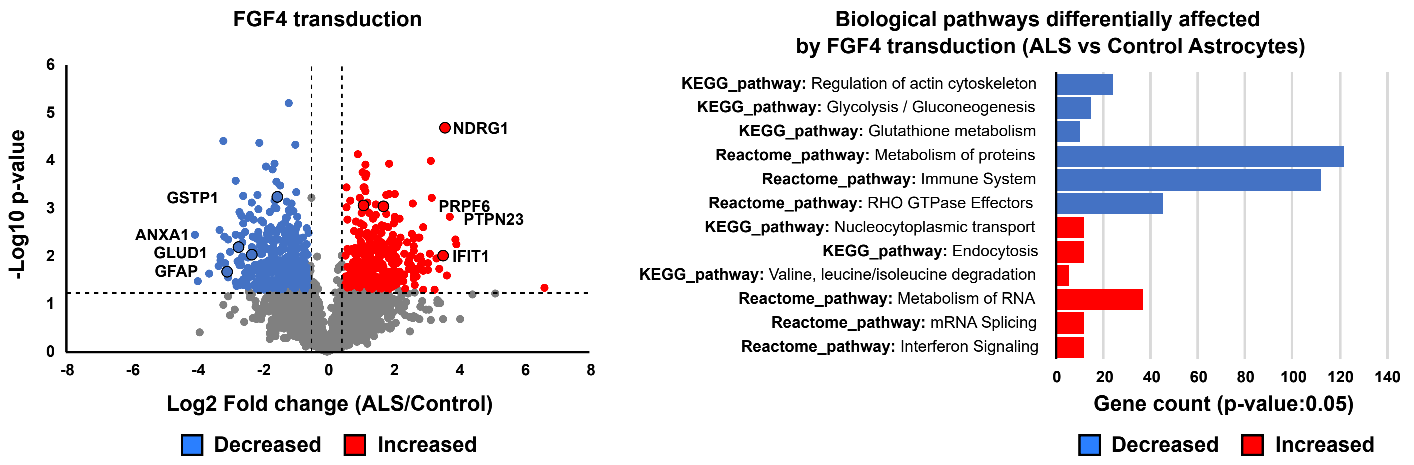

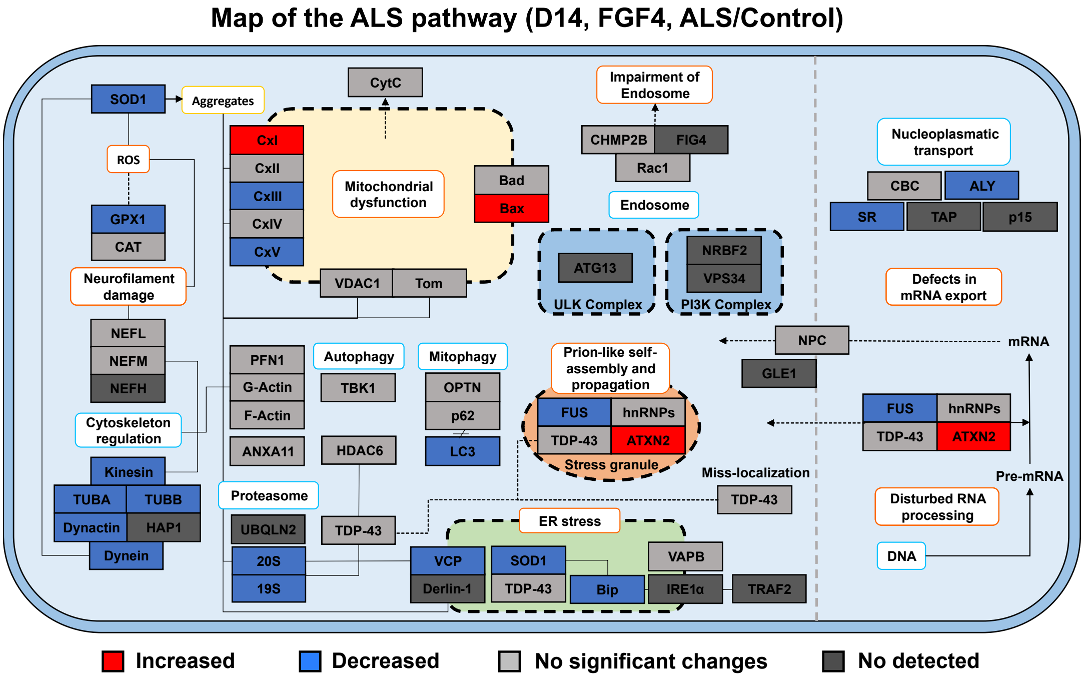

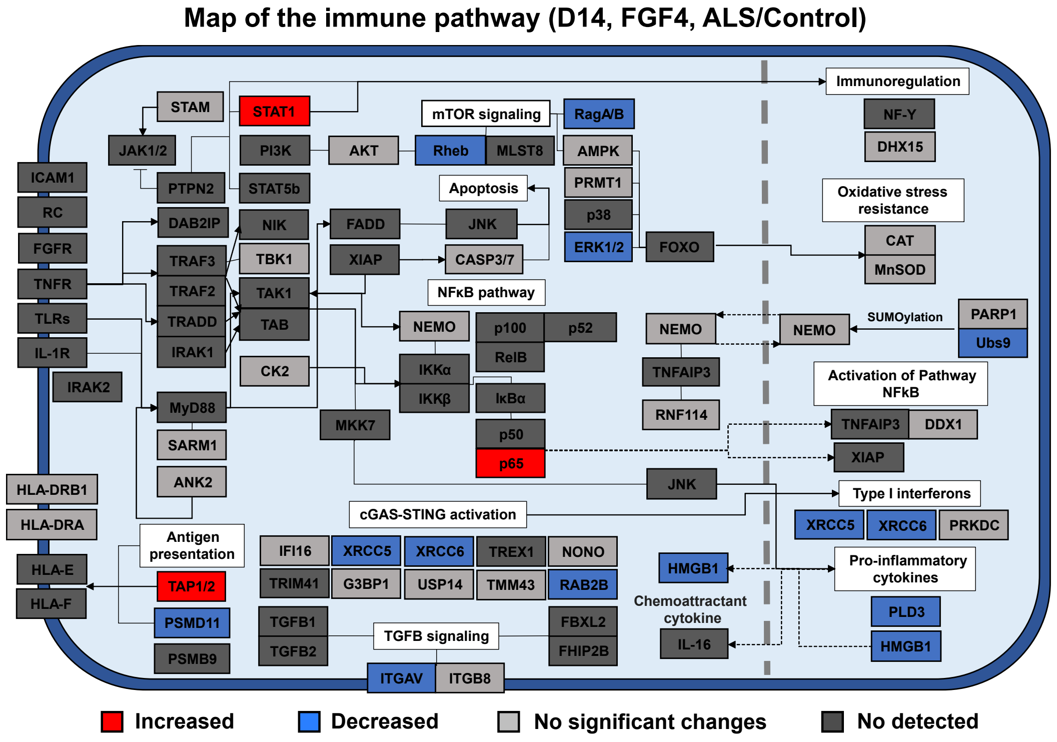

### figure S5

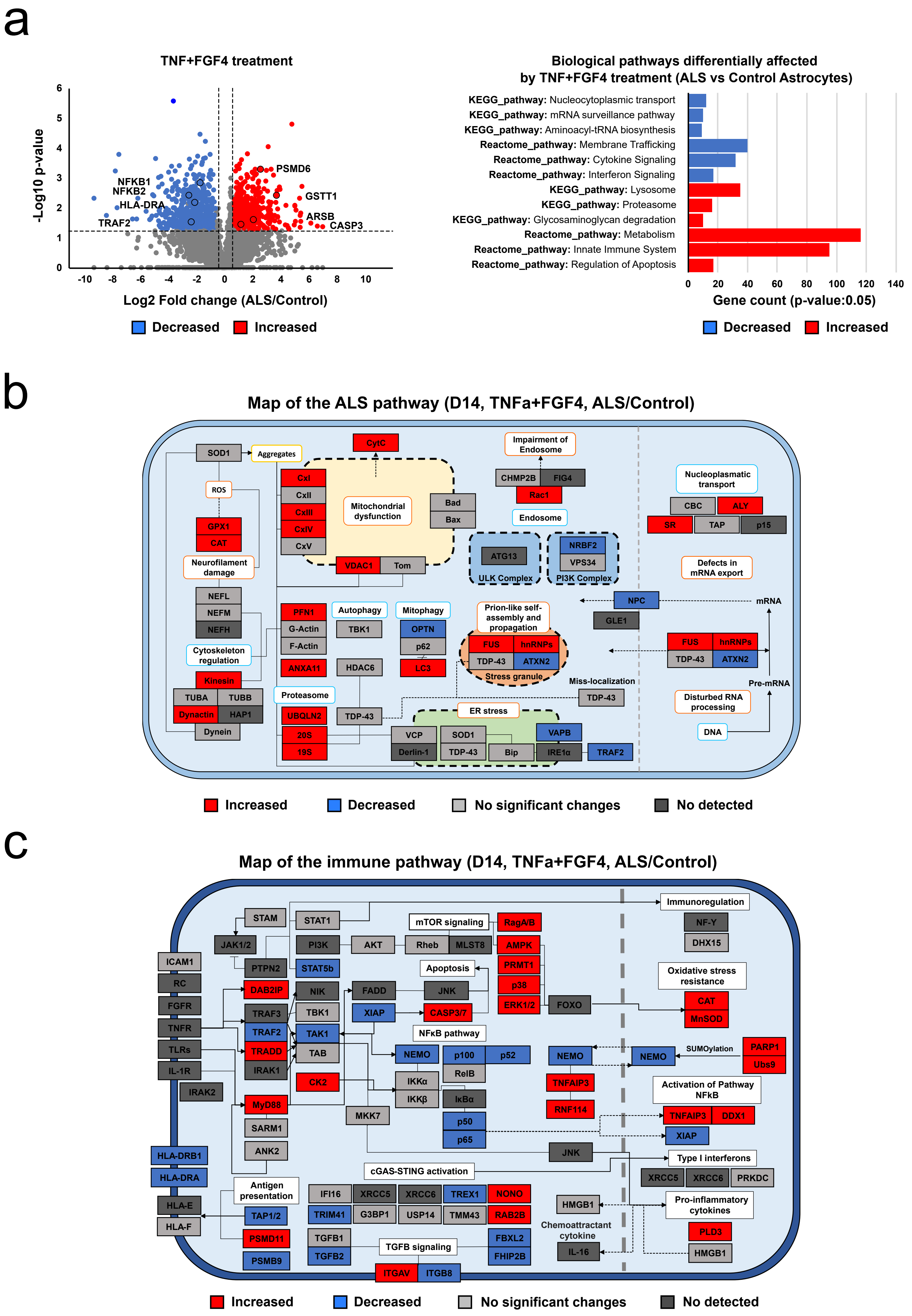
