## Supplementary material for "TNFα hinders FGF4 efficacy to mitigate ALS astrocyte dysfunction and cGAS-STING pathway-induced innate immune reactivity": table S3

**Supplementary Table 3. Summary of the iPSC lines used in this study**

| Diagnosis | Molecular analysis | Age at biopsy | Gender | iPSC line ID | Reference |
|---|---|---|---|---|---|
| Ctrl | NA | 49 | Female | FA11 | TALSCTRL15.12 (RRID:CVCL_FA00) |
| Ctrl | NA | 56 | Female | 37N | Pomeshchik et al., 2020 |
| ALS | SOD1; p.Ala5Val | 65 | Female | NN3854 | ND35671 (RRID:CVCL_T865) |
| ALS | SOD1, p.Ala5Val | 58 | Female | ND35659 | ND35659 (RRID:CVCL_T848) |
| ALS | SOD1; p.Ala5Val | 45 | Female | ND 35673 | ND35673 (RRID:CVCL_Y805) |
